# The China Brain Multi-omics Atlas Project (CBMAP)

**DOI:** 10.1101/2025.04.30.651445

**Authors:** Dan Zhou, Yuan Zhou, Zeyu Sun, Feiyang Ji, Dandan Zhang, Quan Wang, Yijun Ruan, Yongcheng Wang, Yimin Zhu, Xiaohui Sun, Mulin Jun Li, Changzheng Yuan, Kefu Liu, Lingyun Sun, Wenli Zhai, Jiayao Fan, Keqing Zhu, Wenying Qiu, Xiaoxin Yan, Chao Ma, Yi Shen, Aimin Bao, Weihua Yue, Yongyong Shi, Chao Chen, Jian Yang, Shumin Duan, Jing Zhang

**Affiliations:** School of Public Health, the Second Affiliated Hospital, Zhejiang University School of Medicine, Hangzhou, China; National Health and Disease Human Brain Tissue Resource Center, Zhejiang University School of Medicine, Hangzhou, China; Zhejiang Key Laboratory of Intelligent Preventive Medicine, Hangzhou, China; State Key Laboratory for Diagnosis and Treatment of Infectious Diseases, Collaborative Innovation Center for Diagnosis and Treatment of Infectious Diseases, The First Affiliated Hospital, School of Medicine, Zhejiang University, Hangzhou, China; Jinan Microecological Biomedicine Shandong Laboratory, Jinan, China; Department of Surgical Oncology, Sir Run Run Shaw Hospital, Zhejiang University School of Medicine, Hangzhou, China; Provincial Clinical Research Center for CANCER, Hangzhou, China; Department of Pathology, and Department of Medical Oncology of the Second Affiliated Hospital, Zhejiang University School of Medicine, Hangzhou, China; Key Laboratory of Disease Proteomics of Zhejiang Province, Zhejiang University School of Medicine, Hangzhou, China; Department of Biomedical Engineering, College of Automation Engineering, Nanjing University of Aeronautics and Astronautics, Nanjing, China; Life Sciences Institute, Zhejiang University, Hangzhou, China; Liangzhu Laboratory, Zhejiang University, Hangzhou, China; Department of Laboratory Medicine, the First Affiliated Hospital, Zhejiang University School of Medicine, Hangzhou, China; Department of Epidemiology and Biostatistics, and Department of Respiratory Disease, Sir Run Run Shaw Hospital, Zhejiang University School of Medicine, Hangzhou, China; School of Public Health, Zhejiang Chinese Medical University, Hangzhou, China; Guangzhou Women and Children’s Medical Center, Guangzhou Medical University, Guangzhou, China; Center for Medical Genetics & Hunan Key Laboratory of Medical Genetics, School of Life Sciences, and Department of Psychiatry, The Second Xiangya Hospital, Central South University, Changsha, China; Department of Neurology in Second Affiliated Hospital, Key Laboratory of Medical Neurobiology of Zhejiang Province, and Department of Neurobiology, School of Medicine, Zhejiang University, Hangzhou, China; Institute of Basic Medical Sciences Chinese Academy of Medical Sciences, School of Basic Medicine Peking Union Medical College, China; Department of Anatomy & Neurobiology, Central South University Xiangya School of Medicine, Changsha, China; School of Brain Science and Brain Medicine, Zhejiang University School of Medicine, Hangzhou, China; Peking University Sixth Hospital, Peking University Institute of Mental Health, Beijing, China; Institute of Neuroscience, State Key Laboratory of Neuroscience, Center for Excellence in Brain Science and Intelligence Technology (CEBSIT), Chinese Academy of Sciences, Shanghai, China; Bio-X Institutes, Key Laboratory for the Genetics of Developmental and Neuropsychiatric Disorders (Ministry of Education), Shanghai Jiao Tong University, Shanghai, China; MOE Key Laboratory of Rare Pediatric Diseases & Hunan Key Laboratory of Medical Genetics, School of Life Sciences, and Department of Psychiatry, The Second Xiangya Hospital, Central South University, Changsha, China; Hunan Key Laboratory of Animal Models for Human Diseases, Central South University, Changsha, China; School of Life Sciences, Westlake University, Hangzhou, China; Westlake Laboratory of Life Sciences and Biomedicine, Hangzhou, China; Department of Neurology of Second Affiliated Hospital and School of Brain Science and Brain Medicine, Zhejiang University School of Medicine, Hangzhou, China; Liangzhu Laboratory, MOE Frontier Science Center for Brain Science and Brain-Machine Integration, State Key Laboratory of Brain-Machine Intelligence, Zhejiang University, Hangzhou, China; Department of Pathology, The First Affiliated Hospital, Zhejiang University School of Medicine, Hangzhou, China

## Abstract

The China Brain Multi-omics Atlas Project (CBMAP) aims to generate a comprehensive molecular reference map of over 1,000 human brains (Phase I), spanning a broad age range and multiple regions in China, to address the underrepresentation of East Asian populations in brain research. By integrating genome, epigenome, transcriptome, proteome (including multiple post-translational modifications), and metabolome data, CBMAP is set to provide a rich and invaluable resource for investigating the molecular underpinnings of aging-related brain phenotypes and neuropsychiatric disorders. Leveraging high-throughput omics data and advanced technologies, such as spatial transcriptomics, proteomics, and single-nucleus 3D chromatin structure analysis, this atlas will serve as a crucial resource for the brain science community, illuminating disease mechanisms and enhancing the utility of data from genome-wide association studies (GWAS). CBMAP is also poised to accelerate drug discovery and precision medicine for brain disorders.

## Rationale

The onset and progression of neurological and psychiatric disorders are intricately tied to the molecular processes within brain tissues. Research has demonstrated that the levels of transcription and protein, irrespective of the presence of post-translational modifications (PTMs), exhibit a high degree of specificity in brain tissues, making it challenging to use peripheral tissues as a proxy^1, 2^. Non-primates and non-human primates have been used to study the physiology and pathophysiology of the human brain. While the findings have been informative, significant gaps remain in accurately simulating the human brain, particularly for high-level cognitive functions such as self-cognition and theory of mind^3-5^. Furthermore, model animals often exhibit limited interspecies conservation in intergenic regions^3, 6^, restricting the understanding of noncoding regulations. Therefore, a human-based molecular atlas that is specific to brain tissues or cell types is indispensable^2^.

Population-level molecular atlases for humans, such as the 1000 Genomes, have greatly propelled biomedical research by providing a deep catalog of genetic diversity across global populations^7-9^. However, the biology of regulation is missing when solely establishing genetic variant-disease associations, without considering regulatory molecules. A population-level multi-omics reference panel would be fundamental for elucidating the distribution and diversity of molecules, their interrelationships, and their associations with aging and various diseases^1, 10-12^. A multi-omics reference panel for human brain tissues was lacking until recent efforts through projects, such as GTEx, ROSMAP, PsychENCODE, MSBB, and the study by the Knight-ADRC^1, 3, 13-15^. The GTEx project has constructed a joint genome-transcriptome atlas, showing transcriptional distribution across 50 human tissues and revealing the effects of genetic variation on transcriptional regulation^1^. This has facilitated the discovery of potential regulatory molecules that mediate the effects of variants on diseases, as identified from genome-wide association studies (GWAS), thus partially addressing the “missing biology” between variants and diseases^1^. However, the sample sizes for brain tissues in GTEx are limited, with only 100 to 200 sample, most of which are predominantly from individuals of European ancestry^1^. The Religious Orders Study/Memory and Aging Project (ROSMAP) has analyzed the genomes, epigenomes, transcriptomes, and proteomes of hundreds of human brain tissue samples, primarily from individuals of European ancestry, thereby creating a multi-omics atlas mainly focused on understanding the etiology of Alzheimer’s disease (AD)^10, 11, 13, 16-18^. The PsychENCODE project aims to uncover the genetic and molecular mechanisms of psychiatric disorders through omics data from brain tissues, yet it has a limited representation of Asian samples^19-22^. The Mount Sinai/JJ Peters VA Medical Center Brain Bank (MSBB – Mount Sinai NIH Neurobiobank) integrates multiple omics data across various brain regions alongside quantitative measures of neuritic plaque density and clinical dementia ratings to advance understanding of Alzheimer’s disease pathogenesis^15^. Our companion study has suggested that the molecular mechanisms underlying neuropsychiatric disorders might differ between Asian and European populations^23^. It also underscored that increasing genetic ancestral diversity is more efficient for power improvement for probing trait-associated genetic elements than increasing the sample size within single-ancestry reference panel^23^. Therefore, there is an urgent need to fill the gap in large-scale multi-omics brain atlases for Asian populations.

Due to the unique nature of brain tissues, obtaining population-scale samples through hospital- or community-based recruitment is very challenging. National brain banks offer a great opportunity for enabling systematic profiling of the human brain molecular features^24^. Led by the National Health and Disease Human Brain Tissue Resource Center and in collaboration with multiple centers within the China Human Brain Bank Consortium, we propose the China Brain Multi-omics Atlas Project (CBMAP). In Phase I of the project, we collected brain tissue samples from over 1,000 Chinese donors. This project will provide a comprehensive landscape for the molecular profiles across the genome, epigenome, transcriptome, proteome, and metabolome for human brain samples (**Figure 1**). It is worth mentioning that CBMAP will also encompass multiple PTMs that have not been captured in any of the previous studies at the population scale. PTMs are critical mechanisms of protein functional regulation, altering protein properties, localization, stability, and interactions through the addition or removal of specific chemical groups, or through protein cleavage ^25, 26^. These modifications play essential roles in maintaining dynamic cellular functions and biological processes. Changes in the phosphorylation of tau protein are among the most characteristic manifestations of AD ^27^. In addition, we will generate spatial omics and single-nucleus 3D structure of chromatin data for selected samples to provide a higher resolution and deeper insights into the molecular underpinnings of brain-related disorders. The 3D structure of chromatin is critical for gene regulation and cellular function. Chromatin regulates precise gene expression through topologically associating domains (TADs) and spatial interactions between enhancers and promoters ^28^. These mechanisms are especially important in the complex development and function of the brain. Given the currently limited research on brain PTMs and 3D chromatin structures, unknown mechanisms could be revealed for the etiology and progression of brain diseases by a comprehensive study.

**Figure 1.**
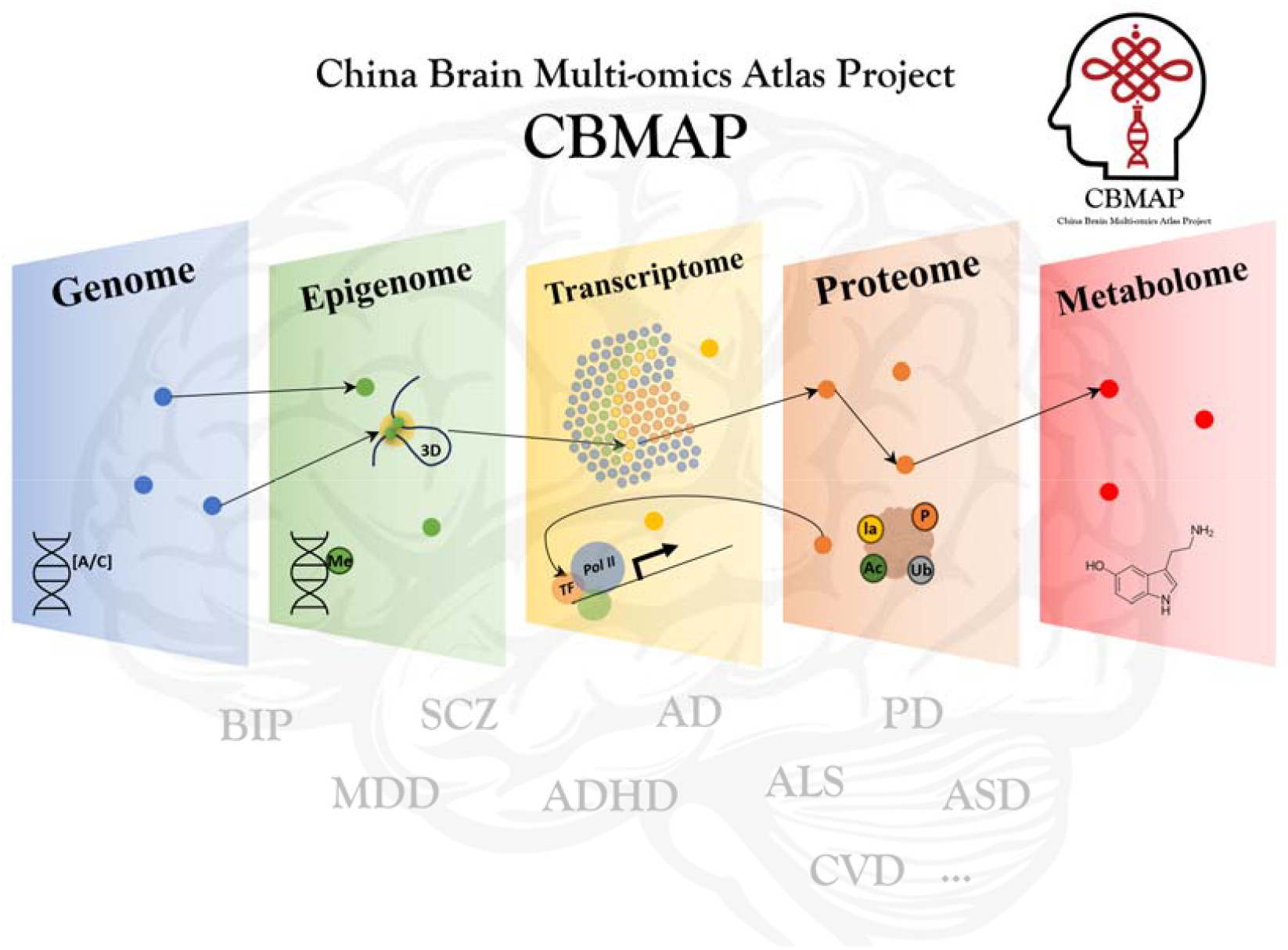
An overview of the China Brain Multi-omics Atlas Project (CBMAP). The project aims to build a multi-omics atlas for the genome, epigenome, transcriptome, proteome, and metabolome. We will study the etiology of multiple brain-related disorders, mainly including Alzheimer’s disease (AD), Parkinson’s disease (PD), cerebrovascular disease (CVD), amyotrophic lateral sclerosis (ALS), schizophrenia (SCZ). The logo in the upper right corner is composed of a Chinese knot and a DNA double helix structure, which resemble the shapes of brain tissue and blood vessels. Chinese knots are more than just handicrafts; they also symbolics of traditional Chinese culture, representing people’s wishes for happiness, safety, health, longevity, and love.

CBMAP, an essential component of the China Brain Project (CBP) and funded in part by CBP, focuses on the aspect of basic research on neural mechanisms underlying cognition ^5^. CBMAP is building a multi-omics reference map of over 1,000 human brains (Phase I), aimed at understanding the molecular networks and features of cognition, aging, and brain-related disorders based on the population level profile.

## Goals of the project

1. To characterize various molecular elements in Chinese population and highlight potential differences within the Chinese population, as well as between Chinese and other populations.
2. To establish a comprehensive atlas of molecules and co-regulation modules associated with aging and neuropsychiatric disorders (including comorbidities) across multiple layers. To investigate potential molecular subtypes for brain related diseases and clinical manifestations.
3. To generate integrated maps of spatial omics, single-nucleus 3D epigenetic interactions, and transcriptional regulation for representative samples, providing deep insights into cellular features underlying brain related conditions.
4. To profile multiple PTMs, uncovering protein-level crosstalks at the population level that have not yet been extensively studied in the European population. This also includes identifying PTM quantitative trait loci (ptmQTLs) and elucidating their local and distal regulatory mechanisms.
5. To elucidate the genetic architecture of various molecules, which may exhibit ancestry-specific, disease-specific, and age-specific characteristics. With a comprehensive spectrum of omics, we aim to estimate the configurations^29^ and potential causal mechanisms of regulatory cascades initialized by genetic variants^30^.
6. Under the hypothesis of “different ancestries, same pathogenic genetic element(s)”, we aim to enhance mapping resolution in the study of disease-associated molecules by integrating multi-ancestry data^31-33^.

## Methods

### Sample and tissue collection

As illustrated in **Figure 2**, Phase I of the CBMAP was initiated by the China Human Brain Bank Consortium following a standardized operational protocol for human brain banking ^34-36^. Members of the consortium including the National Health and Disease Human Brain Tissue Resource Center at Zhejiang University (ZJU) in southeastern China, the National Human Brain Bank for Development and Function at Peking Union Medical College (PUMC) in northern China, and the Xiangya Medical School Brain Bank at Central South University (CSU) in central China take part in the Phase I of the CBMAP, contributing over 1,000 donors in total. These three brain banks are among the earliest established and currently hold the top-ranked brain sample collections in China. The Phase I collection covers the Yangtze River Delta urban cluster by ZJU in the southeast, the Beijing-Tianjin-Hebei metropolitan area by PUMC in the north, and the Central Yangtze River region by CSU in the middle, encompassing populations of 111 million, 245 million, and 110 million, respectively, for a total population coverage of approximately 466 million people (2022 China Urban Development Potential Ranking).

**Figure 2.**
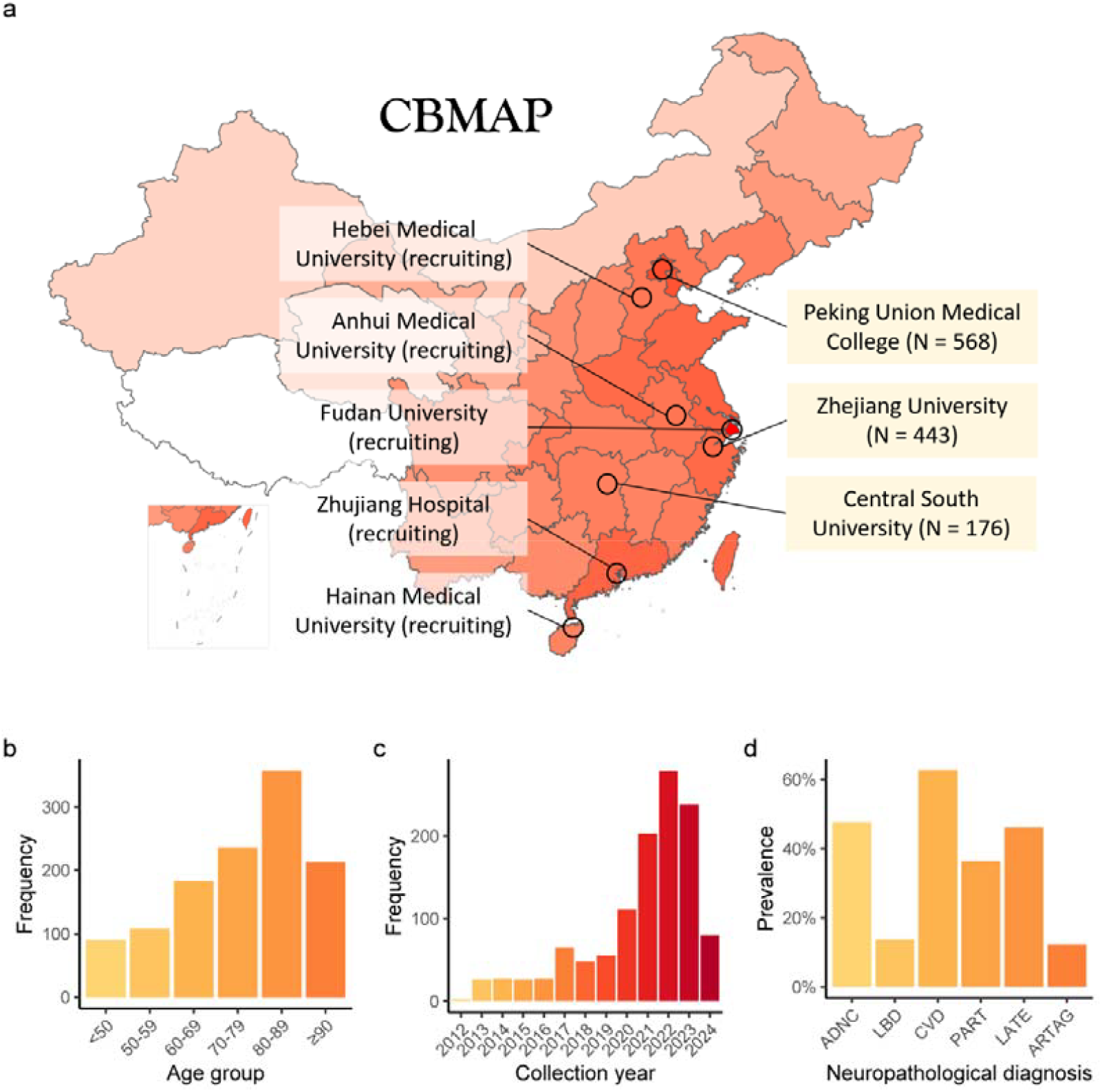
Sample distribution. Brain tissue samples for the CBMAP were collected from multiple brain banks from the China Human Brain Bank Consortium. Phase I of the CBMAP includes the brain banks of Zhejiang University, Peking Union Medical College, and Central South University. Recruitment is ongoing for the remaining brain banks. As shown in panel (a), a deeper red indicates a higher population density. These brain banks cover most of China’s densely populated areas. The age distribution, collection year distribution, and neuropathological diagnosis distribution of the Phase I samples are shown in panels (b), (c), and (d), respectively. ADNC: Alzheimer’s disease neuropathological change, LBD: Lewy body disease, CVD: cerebrovascular diseases, PART: primary age-related tauopathy, LATE: limbic-predominant age-related TDP-43 encephalopathy, ARTAG: aging-related tau astrogliopathy.

All donors or their families provided informed consent through voluntary donation agreements, permitting the use of their information and biospecimens for future studies. The study protocol was approved by the Ethics Committee of Zhejiang University School of Medicine (2020-005 and 2024-007). During sample collection, we recorded the donor’s age, sex, time of death, and disease history. This project only included samples with an ischemia time of less than 24 hours and excluded samples with highly pathogenic infections such as HIV. Donors under 18 years of age were not included in this project. Fetal and child donors will be incorporated in a future extension of the CBMAP, focusing on developmental processes. In Phase II, the sample size will be expanded to approximately 2,000 donors, by incorporating contributions from members of the China Human Brain Bank Consortium, including the Hebei Medical University Brain Bank in northern China, the Fudan University Brain Bank, and the Anhui Medical University Brain Bank in eastern China^37^. We are also actively working to integrate brain banks from the southwestern and northwestern regions of China as well as remote areas, e.g. Hainan, into the project to ensure comprehensive representation of most, if not all, populations in China.

### Brain regions

In Phase I of the CBMAP, we prioritized the region at the intersection of Brodmann Area 9 (BA9) and the superior frontal gyrus. This region primarily overlaps with the dorsolateral prefrontal cortex (DLPFC), which is instrumental in managing various cognitive processes, including working memory, cognitive flexibility, and planning^38^. The broad functional spectrum of DLPFC implicates it in neurodegenerative diseases such as Alzheimer’s disease and a range of psychiatric disorders^3^. The DLPFC has been a focal point for several large-scale projects, including ROSMAP, PsychENOCDE, and GTEx, have also considered this brain region as a primary focus of their research^1, 3, 17^. In subsequent phases of the CBMAP, we will extend our research to encompass additional brain regions including hippocampus, striatum, amygdala, substantia nigra, and hypothalamus, thereby constructing a more comprehensive molecular atlas.

### Neuropathological diagnosis

Samples collected in this project come with definitive pathological diagnoses, including primary age-related tauopathy (PART)^39^, limbic-predominant age-related TDP-43 encephalopathy (LATE)^40^, aging-related tau astrogliopathy (ARTAG)^41^, Alzheimer’s disease neuropathological change (ADNC)^42^, Lewy body disease (LBD)^43^, cerebrovascular diseases (CVD)^44^, and pathologically healthy controls. These uniformly applied pathological diagnostic procedures are organized by the National Health and Disease Human Brain Tissue Resource Center at Zhejiang University, following a unified process of quality controls as detailed in the Supplementary materials and our recent comorbidity study^37^. The molecular features of pathological conditions and the pathological condition-specific regulation patterns (including genetic regulations) will be studied.

### Genomics and epigenomics

We will conduct whole-genome sequencing (WGS). Considering factors such as sample size, frequency of rare variants, and sequencing costs, the sequencing depth for Phase I is set at between 10-20×. DNA methylation will be assessed using the Infinium MethylationEPIC v2.0 BeadChip. Additionally, we will carry out single-nucleus ATAC sequencing (snATAC-seq) on selected samples to capture cell-type-specific chromatin accessibility. Techniques such as ATAC-Me, which combine DNA methylation and chromatin accessibility measurements, will be considered to gain deeper insights into their joint regulation of gene expression^45^. To explore interactions among regulatory elements, we plan to conduct capture-Hi-C^46^. Moreover, we will employ a single-cell/nucleus tri-omic approach ChAIR (chromatin accessibility, interaction, and RNA simultaneously) for single-cell 3D epigenomic and transcriptional regulatory panoramic scanning^47^.

### Transcriptomics

To profile the transcriptome, we will conduct ribo-free bulk RNA-seq to cover as many samples as possible with RIN ≥ 5. In addition, we will conduct single-nucleus RNA sequencing (snRNA-seq) for selected samples to profile cell-type-specific regulations and to support the deconvolution-based cell-type proportion and cell-type level gene expression estimation. Random primers-based techniques including snRandom-seq will be implemented to capture the low-quality RNAs^48^. Long-read RNA-seq will be utilized to achieve a higher quality of identification of splicing events^49, 50^. Furthermore, we are conducting spatial transcriptomics on representative samples and regions to study the etiology of neurodegenerative diseases^51^.

The representative samples are selected based on research questions. To probe disease-related transcriptomic features, we will select cases with clear pathological/clinical diagnoses (including different disease stages and comorbidities) and control samples matched by potential confounding factors. For identifying the molecular features of natural physiological processes (e.g., aging), we will select samples based on the distribution of age, sex, and region. These two parts will cover 100-200 samples of prefrontal cortex.

Proteomics, post-translational modifications, and metabolomics Protein and post-translational modification (PTM) abundance will be measured using Liquid Chromatography-Mass Spectrometry (LC-MS). This will enable us to investigate protein and PTM profiles of serine/threonine phosphorylation, ubiquitination, and acetylation modifications. Additionally, we plan to examine other PTMs including lactylation to understand the hypoxia-related molecular consequences. For the samples planned for spatial transcriptomics and single-nucleus transcriptomics, we will also conduct spatial proteomics and PTM (at least for phosphorylomics) measurements to study the potential molecular functions within specific microenvironments across multiple omics layers.

In addition, we plan to perform metabolomics analysis on brain tissue homogenates and high-resolution spatial metabolomics. Homogenate metabolomics will leverage widely targeted LC-MS technology, complemented by targeted metabolomics for verification.

### Data visualization

We will develop a data visualization portal to facilitate access by the scientific community. The distribution of basic information including age, sex, and sample source will be visually presented on the website. Spatial omics images of several key brain regions will also be available on the website. Results generated from the analysis, such as genetic-molecular trait association signals (i.e., xQTLs), genetic-informed molecular trait-disease association results, and case-control differential analysis for neuropsychiatric disorders, will be made available on the portal, with interactive query and download functionalities.

## The current stage overview

In Phase I, we collected tissue samples from 1,187 Chinese donors from the brain banks at Zhejiang University (N = 443), Peking Union Medical College (N = 568), and Central South University (N = 176). In the pilot stage of Phase I, we have completed the bulk-level measurement for the genome, epigenome, transcriptome, proteome, and phosphoproteome profile in a subset of samples (**Figure S1**) and established the data processing and quality control pipeline. High-throughput data generation is underway for all available samples. In addition, we are generating snRNA-seq data, snATAC-seq data, and single nucleus-level 3D epigenomic and transcriptional regulatory maps for representative tissue samples, with particular interest in neurodegenerative diseases including AD. Moreover, we have finished pathological diagnoses including PART, LATE, ARTAG, ADNC, LBD, and CVD for most samples.

As shown in **Figure 2a**, these samples represent the most densely populated regions in the northern, southeastern, and central parts of China. Among the donors, 37.4% are females, with a median age of 78 years and an interquartile range (IQR) of 66 (P_25_) to 87 (P_75_) years (**Figure 2b)**. Since 2012, the number of donors has generally increased year by year (**Figure 2c**). The detection rates of age-related neuropathological changes such as PART, LATE, and ARTAG among donors are 36.4%, 46.1%, and 12.2%, respectively (**Figure 2d**). The detection rates for ADNC, LBD, and CVD are 47.6%, 13.7%, and 62.6% respectively (**Figure 2d**). In addition, 24 donors, aged 27 to 96, have clinical records with schizophrenia diagnosis. We are also enhancing access to de-identified healthcare records, including high-resolution brain MRI imaging, clinical diagnoses, and laboratory measurements.

In addition, we evaluated the potential impact of sample quality-related characteristics (including RIN and PMI) on molecular profiles based on the pilot data. We performed principal component analysis and presented the distribution of RIN (**Figures S1a-S1d**) and PMI (**Figures S1e-S1h**) on the top PCs of molecular traits including epigenome (DNA methylation), transcriptome, proteome, and phosphoproteome. In line with the observation in the GTEx v8 prefrontal cortex BA9 samples (**Figure S2**), the RNA degradation level can be captured by PC1 of the gene expression profile. No clear patterns emerged between the other molecular profiles and sample quality characteristics. These preliminary results suggest that gene expression profile is sensitive to RIN, while the PMI (within 24 hours) plays a limited role in these molecular profiles. RIN, PMI, age, and sex will be adjusted as covariates for regression models. Compared to ROSMAP, preliminary cis-eQTL analysis in CBMAP samples revealed a greater number of SNP-gene expression associations with an FDR < 0.05 (**Figure S3**). The greater number of cis-eQTLs may be attributed to a larger sample size of CBMAP than ROSMAP. A concordant trans-eQTL pattern was observed between CBMAP and ROSMAP (**Figure S3**). The observations above indicated a good tissue sample quality.

## Comparison with other projects

Compared with the GTEx (we only refer to brain tissues in this context), ROSMAP, PsychENCODE, MSBB, Banner, and the project by Knight-ADRC^1, 3, 13, 14, 52^, the CBMAP stands out in several key aspects (**Table S1**): (1) we have included over 1,000 donors in Phase I and will increase the sample size to 2,000 in the next phase; (2) all the samples are Chinese, a population scarcely represented in previous studies; (3) alongside single-nucleus sequencing, bulk-level high-throughput measurements will be systematically conducted across the entire molecular spectrum, from the genome to the metabolome; (4) high-resolution spatial omics will be profiled for representative samples; (5) single nucleus-level 3D genome interaction will be revealed; (6) multiple post-translational modifications will be profiled and associated with other molecular layers, population characteristics, and diseases.

We have also compared the sample characteristics across the projects. In Phase I, we took samples from the BA9 region and conducted RIN value measurement, which showed that the median and interquartile range of RIN values were comparable to those of the GTEx project’s prefrontal cortex (PFC) BA9 region samples and the ROSMAP project’s PFC region samples (**Figure 3a**). The median and interquartile range of postmortem interval (PMI) for CBMAP samples were lower than those of the GTEx project and comparable to those of the ROSMAP project (**Figure 3b**). Notably, CBMAP covers a much broader age range than either GTEx or ROSMAP (**Figure 3c**). Only minimal differences were observed across CBMAP centers (**Figures 3d, 3e**, and **3f**). We also note that the proportion of female donors in the CBMAP project is 37.4%. Similarly, in the PsychENCODE project, the proportion of female samples in the control group is also below 40%^53^. The reasons behind these differences warrant further investigation.

**Figure 3.**
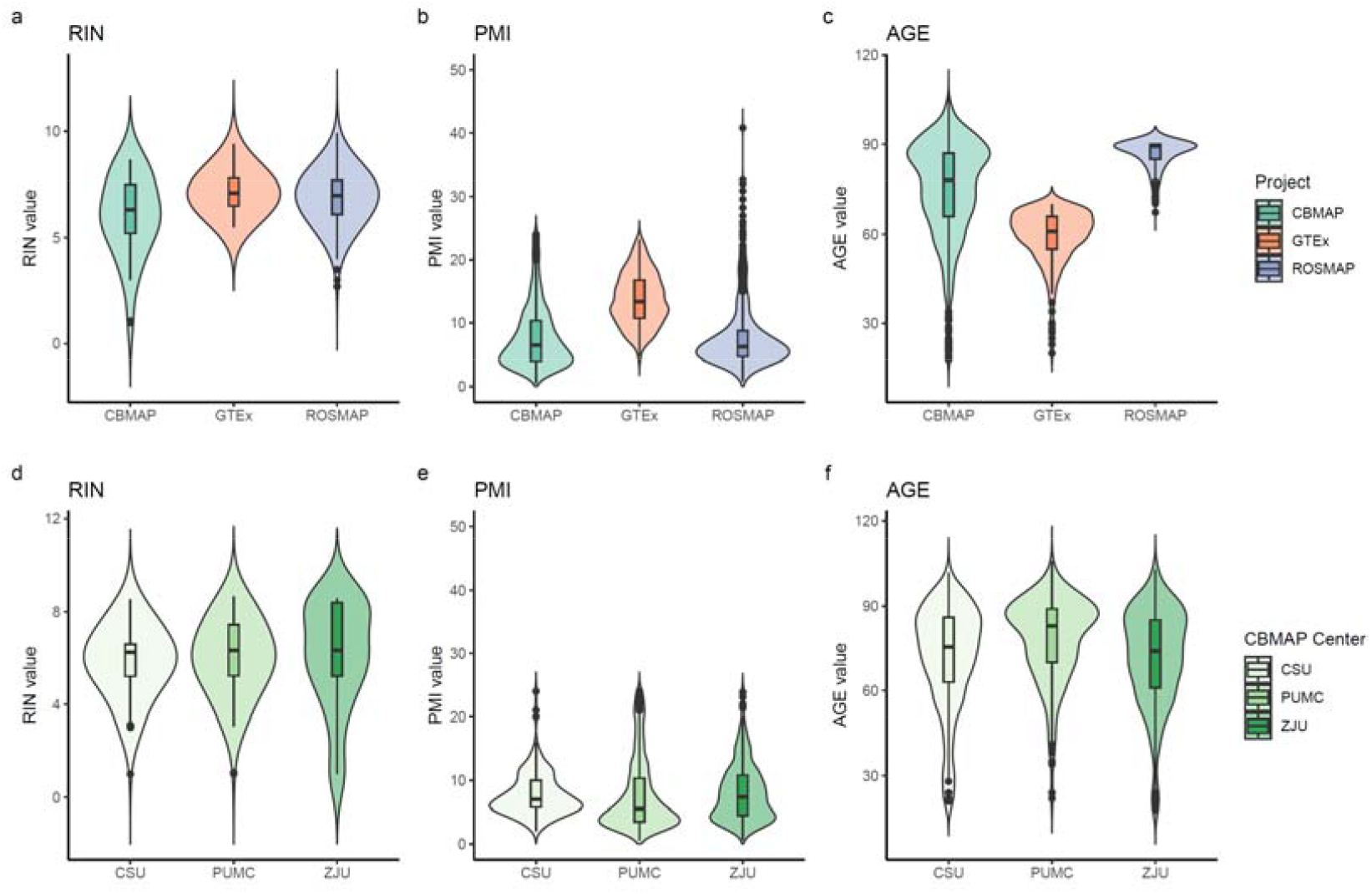
Distribution of RNA integrity, postmortem interval (PMI), and sample age across different datasets. Panels (a), (b), and (c) show the RIN values, PMI (hours), and age for the CBMAP, GTEx (prefrontal cortex BA9), and ROSMAP projects, respectively. Panels (d), (e), and (f) display the distribution of RIN, PMI, and age for the three centers in the CBMAP Phase I. The violin plots illustrate the distribution characteristics of the data. The box spans the interquartile range (IQR), with the median indicated by a horizontal line inside each box. Whiskers extend to the smallest and largest values within 1.5 times the IQR. Outliers beyond this range are plotted as individual points. CSU: Central South University; PUMC: Peking Union Medical College; ZJU: Zhejiang University.

Comparisons will be made for the regulation of molecules across different projects and different ancestries. Meanwhile, variant- and gene-level fine-mapping will be facilitated to have a better resolution by leveraging multi-ancestry datasets.

## Prospectives

### The power of genetics

The completion of the Asian reference panel will enable the bridging of SNPs and diseases by GWAS-based transcriptome-wide association study (TWAS), proteome-wide association study (PWAS), and other genetics-informed studies^54^. The availability of data from Asian populations will facilitate cross-ancestry fine-mapping, enabling more accurate identification of trait-associated molecules^31, 33, 54^. Unlike case-control studies, which can be biased by confounding factors and reverse causation, research based on germline variants offers a natural advantage in causal inference, making it a powerful method for identifying pathogenic molecules. Such genetics-based probing provides crucial support for drug development and repurposing. Recent studies have shown that the success rate of drug development backed by genetic evidence is 2.6 times greater than that without genetic evidence^55^, offering a strong basis for future research leveraging the power of genetics^56^.

### A live cohort of potential donors

The project is supported by the National Health and Disease Human Brain Tissue Resource Center, which is actively constructing a living cohort for potential donors. This setting will enable more accurate pre-mortem data collection including assessments of cognitive functions, lifestyle preferences, environmental factors, and laboratory measurements. Such comprehensive data collection will be invaluable for future research, particularly in exploring gene-environment interactions^5^. Additionally, this initiative enables the collection of peripheral samples, including blood, cerebrospinal fluid, and skin fibroblasts, which can be used to generate induced pluripotent stem cells (iPSCs). These iPSCs will be used to create *in vitro* models that mimic human conditions, facilitating the following validation^5^.

### CBMAP Phase II

In Phase II of CBMAP, we aim to significantly expand and diversify our resources to provide deeper insights into the molecular landscape of the human brain. Building on the foundation of Phase I, we plan to increase the atlas’s sample size from ∼1,000 to ∼2,000. Additional brain banks from the Hebei, Anhui, Shanghai, and Guangzhou, as well as brain banks from more remote areas, such as Hainan, will be included to ensure comprehensive representation of most, if not all, populations in China. During Phase II, we will specifically gather brain samples from donors under 18 and fetal brain samples. Multi-omics analysis will be conducted on 300-500 samples from key brain regions including hippocampus, striatum, amygdala, substantia nigra, and hypothalamus, with a focus on several major neurological diseases. Additionally, 5-10 samples will undergo whole-brain spatial transcriptomics and proteomics mapping. This will be complemented by high-resolution MRI-based 3D coordinate mapping for each tissue sample.

We also plan to include living cohorts, such as specialized disease cohorts (e.g., mental disorders and amyotrophic lateral sclerosis [ALS]) and brain-disease-free aging cohorts. We will integrate the health and medical records of these living cohorts, along with molecular characteristics of peripheral samples (e.g., blood and feces), to investigate the connections between peripheral and brain tissues and their relationships with diseases.

The goals for Phase II include establishing a larger and more regionally representative multi-omics atlas of Chinese brain tissue, validating the findings from Phase I, and uncovering new molecular features. We also aim to identify disease-related spatial molecular signatures across multiple brain regions and complete a comprehensive spatial molecular feature map of the entire brain. Importantly, by integrating peripheral tissue data with post-mortem brain samples, we seek to create a network of molecular profiles that bridges living and post-mortem analyses, advancing our understanding of systemic molecular interconnections.

## Project organization and data sharing

The CBMAP is spearheaded by the China National Health and Disease Human Brain Tissue Resource Center, leveraging the China Human Brain Bank Consortium for tissue collection. A centralized hub, supported by specialized cores for sample collection, pathology, high-throughput data analysis, methodology development, data visualization, data management, and sharing, ensures the efficient execution and scalability of the project (**Figure S4**).

For data sharing, the raw sequencing data will be uploaded to the GSA platform of the China National Bioinformatics Center (https://ngdc.cncb.ac.cn/gsa/). Researchers can apply for data use either through the National Health and Disease Human Brain Tissue Resource Center (http://zjubrainbank.zju.edu.cn/index) or via the GSA platform. All applications will be reviewed and approved by the academic committee and the CBMAP project team.

## Conclusion

The CBMAP is pioneering a comprehensive molecular atlas of the human brain, spanning a wide range of ages and standardized pathological diagnoses. By mapping the entire central dogma, including multiple PTMs, the metabolome, spatial omics, and single-nucleus 3D genome structures, we are laying a crucial foundation for unraveling the complex mechanisms underlying brain function. Our progress in this endeavor brings us closer to understanding the mysteries of the human mind and advancing our ability to address brain-related disorders.

## Acknowledgments

We express our gratitude to all the brain tissue donors and pay tribute to them and their families.

We are thankful to Dr. Chunyu Liu, Dr. Nancy J. Cox, Dr. Hailiang Huang, and Dr. Eric R. Gamazon for critical reading of the manuscript and the feedback.

The project is supported by the STI2030-Major Project (Brain Tissue Resource Repository and Brain Bank Collaboration Network Platform, 2021ZD0201100, [J.Z.]), the National Natural Science Foundation of China (82020108012 [J.Z.], 82204118 [D.Z.], 82022024 [C.C.], 32270656 [D.Z.], and 82370612 [Z.S.]) and the Key R&D Program of Zhejiang (2024C03098, J.Z.).

## Code availability

The scripts are available at https://github.com/zdangm/CBMAP_profile.

## Ethics declarations

### Ethics approval

All donors or their families provided informed consent through voluntary donation agreements, permitting the use of their information and biospecimens for future studies. The study protocol was approved by the Ethics Committee of Zhejiang University School of Medicine (2020-005 and 2024-007).

## Reference

1. Consortium G. The GTEx Consortium atlas of genetic regulatory effects across human tissues. Science 2020; 369(6509): 1318–1330.

2. Gamazon ER, Zwinderman AH, Cox NJ, Denys D, Derks EM. Multi-tissue transcriptome analyses identify genetic mechanisms underlying neuropsychiatric traits. Nature genetics 2019; 51(6): 933–940.

3. Akbarian S, Liu C, Knowles JA, Vaccarino FM, Farnham PJ, Crawford GE et al. The psychencode project. Nature neuroscience 2015; 18(12): 1707–1712.

4. Johnson MB, Kawasawa YI, Mason CE, Krsnik Ž, Coppola G, Bogdanović D et al. Functional and evolutionary insights into human brain development through global transcriptome analysis. Neuron 2009; 62(4): 494–509.

5. Poo MM, D. JL, Ip NY, Xiong ZQ, Xu B, Tan T. China Brain Project: Basic Neuroscience, Brain Diseases, and Brain-Inspired Computing. Neuron 2016; 92(3): 591–596.

6. Konopka G, Friedrich T, Davis-Turak J, Winden K, Oldham MC, Gao F et al. Human-specific transcriptional networks in the brain. Neuron 2012; 75(4): 601–617.

7. Gibbs RA. The human genome project changed everything. Nature Reviews Genetics 2020; 21(10): 575–576.

8. Claussnitzer M, Cho JH, Collins R, Cox NJ, Dermitzakis ET, Hurles ME et al. A brief history of human disease genetics. Nature 2020; 577(7789): 179–189.

9. Consortium GP. A global reference for human genetic variation. Nature 2015; 526(7571): 68.

10. Fujita M, Gao Z, Zeng L, McCabe C, White CC, Ng B et al. Cell subtype-specific effects of genetic variation in the Alzheimer’s disease brain. Nature Genetics 2024; 56(4): 605–614.

11. Mathys H, Boix CA, Akay LA, Xia Z, Davila-Velderrain J, Ng AP et al. Single-cell multiregion dissection of Alzheimer’s disease. Nature 2024: 1–11.

12. Johnson EC, Carter EK, Dammer EB, Duong DM, Gerasimov ES, Liu Y et al. Large-scale deep multi-layer analysis of Alzheimer’s disease brain reveals strong proteomic disease-related changes not observed at the RNA level. Nature neuroscience 2022; 25(2): 213–225.

13. Bennett DA, Buchman AS, Boyle PA, Barnes LL, Wilson RS, Schneider JA. Religious orders study and rush memory and aging project. Journal of Alzheimer’s disease 2018; 64(s1): S161–S189.

14. Yang C, Farias FH, Ibanez L, Suhy A, Sadler B, Fernandez MV et al. Genomic atlas of the proteome from brain, CSF and plasma prioritizes proteins implicated in neurological disorders. Nature neuroscience 2021; 24(9): 1302–1312.

15. Wang M, Beckmann ND, Roussos P, Wang E, Zhou X, Wang Q et al. The Mount Sinai cohort of large-scale genomic, transcriptomic and proteomic data in Alzheimer’s disease. Scientific data 2018; 5(1): 1–16.

16. Xiong X, James BT, Boix CA, Park YP, Galani K, Victor MB et al. Epigenomic dissection of Alzheimer’s disease pinpoints causal variants and reveals epigenome erosion. Cell 2023; 186(20): 4422-4437. e4421.

17. De Jager PL, Ma Y, McCabe C, Xu J, Vardarajan BN, Felsky D et al. A multi-omic atlas of the human frontal cortex for aging and Alzheimer’s disease research. Scientific data 2018; 5(1): 1–13.

18. Ng B, White CC, Klein H-U, Sieberts SK, McCabe C, Patrick E et al. An xQTL map integrates the genetic architecture of the human brain’s transcriptome and epigenome. Nature neuroscience 2017; 20(10): 1418–1426.

19. Ruzicka WB, Mohammadi S, Fullard JF, Davila-Velderrain J, Subburaju S, Tso DR et al. Single-cell multi-cohort dissection of the schizophrenia transcriptome. Science 2024; 384(6698): eadg5136.

20. Huuki-Myers LA, Spangler A, Eagles NJ, Montgomery KD, Kwon SH, Guo B et al. A data-driven single-cell and spatial transcriptomic map of the human prefrontal cortex. Science 2024; 384(6698): eadh1938.

21. Wen C, Margolis M, Dai R, Zhang P, Przytycki PF, Vo DD et al. Cross-ancestry atlas of gene, isoform, and splicing regulation in the developing human brain. Science 2024; 384(6698): eadh0829.

22. Wang D, Liu S, Warrell J, Won H, Shi X, Navarro FC et al. Comprehensive functional genomic resource and integrative model for the human brain. Science 2018; 362(6420): eaat8464.

23. Chen Y, Liu S, Ren Z, Wang F, Liang Q, Jiang Y et al. Cross-ancestry analysis of brain QTLs enhances interpretation of schizophrenia genome-wide association studies. The American Journal of Human Genetics.

24. Wang L, Xia Y, Chen Y, Dai R, Qiu W, Meng Q et al. Brain Banks Spur New Frontiers in Neuropsychiatric Research and Strategies for Analysis and Validation. Genomics Proteomics Bioinformatics 2019; 17(4): 402–414.

25. Vidal CJ. Post-translational modifications in health and disease, vol. 13. Springer Science & Business Media 2010.

26. Martin L, Latypova X, Terro F. Post-translational modifications of tau protein: implications for Alzheimer’s disease. Neurochemistry international 2011; 58(4): 458–471.

27. Scheltens P, De Strooper B, Kivipelto M, Holstege H, Chételat G, Teunissen CE et al. Alzheimer’s disease. Lancet 2021; 397(10284): 1577–1590.

28. Beagan JA, Phillips-Cremins JE. On the existence and functionality of topologically associating domains. Nature genetics 2020; 52(1): 8–16.

29. Wu Y, Qi T, Wray NR, Visscher PM, Zeng J, Yang J. Joint analysis of GWAS and multi-omics QTL summary statistics reveals a large fraction of GWAS signals shared with molecular phenotypes. Cell Genom 2023; 3(8): 100344.

30. Li YI, van de Geijn B, Raj A, Knowles DA, Petti AA, Golan D et al. RNA splicing is a primary link between genetic variation and disease. Science 2016; 352(6285): 600–604.

31. Lu Z, Gopalan S, Yuan D, Conti DV, Pasaniuc B, Gusev A et al. Multi-ancestry fine-mapping improves precision to identify causal genes in transcriptome-wide association studies. Am J Hum Genet 2022; 109(8): 1388–1404.

32. Chen F, Wang X, Jang SK, Quach BC, Weissenkampen JD, Khunsriraksakul C et al. Multi-ancestry transcriptome-wide association analyses yield insights into tobacco use biology and drug repurposing. Nat Genet 2023; 55(2): 291–300.

33. Li Z, Zhao W, Shang L, Mosley TH, Kardia SLR, Smith JA et al. METRO: Multi-ancestry transcriptome-wide association studies for powerful gene-trait association detection. Am J Hum Genet 2022; 109(5): 783–801.

34. Qiu W, Zhang H, Bao A, Zhu K, Huang Y, Yan X et al. Standardized operational protocol for human brain banking in China. Neuroscience Bulletin 2019; 35: 270–276.

35. Xue W, Juanli W, Naili W, Di Z, Zhen C, Hanlin Z et al. Standardized Operational Protocol for the China Human Brain Bank Consortium. Human Brain 2022; 1(1): 92–106-192–106.

36. Ma C, Bao A-M, Yan X-X, Swaab DF. Progress in human brain banking in China. vol. 35. Springer 2019, pp 179–182.

37. Wang X, Zhu K, Wu W, Zhou D, Lu H, Du J et al. Prevalence of mixed neuropathologies in age-related neurodegenerative diseases: A community-based autopsy study in China. Alzheimer’s & Dementia 2024.

38. Carlén M. What constitutes the prefrontal cortex? Science 2017; 358(6362): 478–482.

39. Crary JF, Trojanowski JQ, Schneider JA, Abisambra JF, Abner EL, Alafuzoff I et al. Primary age-related tauopathy (PART): a common pathology associated with human aging. Acta Neuropathol 2014; 128(6): 755–766.

40. Nelson PT, Dickson DW, Trojanowski JQ, Jack CR, Boyle PA, Arfanakis K et al. Limbic-predominant age-related TDP-43 encephalopathy (LATE): consensus working group report. Brain 2019; 142(6): 1503–1527.

41. Kovacs GG, Ferrer I, Grinberg LT, Alafuzoff I, Attems J, Budka H et al. Aging-related tau astrogliopathy (ARTAG): harmonized evaluation strategy. Acta Neuropathol 2016; 131(1): 87–102.

42. Montine TJ, Phelps CH, Beach TG, Bigio EH, Cairns NJ, Dickson DW et al. National Institute on Aging-Alzheimer’s Association guidelines for the neuropathologic assessment of Alzheimer’s disease: a practical approach. Acta Neuropathol 2012; 123(1): 1–11.

43. McKeith IG, Boeve BF, Dickson DW, Halliday G, Taylor JP, Weintraub D et al. Diagnosis and management of dementia with Lewy bodies: Fourth consensus report of the DLB Consortium. Neurology 2017; 89(1): 88–100.

44. Skrobot OA, Attems J, Esiri M, Hortobágyi T, Ironside JW, Kalaria RN et al. Vascular cognitive impairment neuropathology guidelines (VCING): the contribution of cerebrovascular pathology to cognitive impairment. Brain 2016; 139(11): 2957–2969.

45. Guerin LN, Barnett KR, Hodges E. Dual detection of chromatin accessibility and DNA methylation using ATAC-Me. Nature protocols 2021; 16(12): 5377–5397.

46. Freire-Pritchett P, Ray-Jones H, Della Rosa M, Eijsbouts CQ, Orchard WR, Wingett SW et al. Detecting chromosomal interactions in Capture Hi-C data with CHiCAGO and companion tools. Nature protocols 2021; 16(9): 4144–4176.

47. Chai H, Huang X, Xiong G, Huang J, Pels KK, Meng L et al. Tri-omic mapping revealed concerted dynamics of 3D epigenome and transcriptome in brain cells. bioRxiv 2024: 2024.2005. 2003.592322.

48. Xu Z, Zhang T, Chen H, Zhu Y, Lv Y, Zhang S et al. High-throughput single nucleus total RNA sequencing of formalin-fixed paraffin-embedded tissues by snRandom-seq. Nature communications 2023; 14(1): 2734.

49. Pardo-Palacios FJ, Wang D, Reese F, Diekhans M, Carbonell-Sala S, Williams B et al. Systematic assessment of long-read RNA-seq methods for transcript identification and quantification. Nature methods 2024: 1–15.

50. Kovaka S, Zimin AV, Pertea GM, Razaghi R, Salzberg SL, Pertea M. Transcriptome assembly from long-read RNA-seq alignments with StringTie2. Genome biology 2019; 20: 1–13.

51. Wei X, Fu S, Li H, Liu Y, Wang S, Feng W et al. Single-cell Stereo-seq reveals induced progenitor cells involved in axolotl brain regeneration. Science 2022; 377(6610): eabp9444.

52. Beach TG, Adler CH, Sue LI, Serrano G, Shill HA, Walker DG et al. Arizona Study of Aging and Neurodegenerative Disorders and Brain and Body Donation Program. Neuropathology 2015; 35(4): 354–389.

53. Xia Y, Xia C, Jiang Y, Chen Y, Zhou J, Dai R et al. Transcriptomic sex differences in postmortem brain samples from patients with psychiatric disorders. Sci Transl Med 2024; 16(749): eadh9974.

54. Gamazon ER, Wheeler HE, Shah KP, Mozaffari SV, Aquino-Michaels K, Carroll RJ et al. A gene-based association method for mapping traits using reference transcriptome data. Nat Genet 2015; 47(9): 1091–1098.

55. Minikel EV, Painter JL, Dong CC, Nelson MR. Refining the impact of genetic evidence on clinical success. Nature 2024; 629(8012): 624–629.

56. Razuvayevskaya O, Lopez I, Dunham I, Ochoa D. Genetic factors associated with reasons for clinical trial stoppage. Nature Genetics 2024: 1–6.

