## supplemental table and figure for "The China Brain Multi-omics Atlas Project (CBMAP)"

**Supplementary** **materials**

**Supplementary texts**

**Neuropathological diagnosis**

As part of the autopsy process, each brain underwent a comprehensive series of evaluations. Initially, the brains were weighed and then fixed in 10% buffered formalin for at least two weeks. After fixation, they were carefully sectioned into coronal slices. Macroscopic abnormalities, including lesions and vascular pathologies, were systematically examined. Standardized tissue samples were then collected from 22 distinct brain regions, as described in Wang et al^1^. Immunohistochemical (IHC) staining was performed, with details of the specific staining protocols provided in Wang et al^1^.

To ensure uniformity and reliability, we applied standardized assessment criteria across all the brain banks in the CBMAP. IHC findings were interpreted following established staging frameworks, including the Thal phasing for Aβ, the Braak stage for hyperphosphorylated tau, the BrainNet Europe classification for α-synuclein, and assessments for TDP-43 pathology.

The extent and severity of Alzheimer's disease neuropathological changes (ADNC) were determined using the “ABC” grading system, which categorizes pathology as low, intermediate, or high^2^. Lewy body disease (LBD), encompassing both Parkinson’s disease (PD) and incidental PD, was diagnosed based on the pathological presence of α-synuclein, even in individuals without clinical PD symptoms^3^. A comprehensive evaluation of vascular pathologies was conducted, encompassing macroscopic and microscopic infarcts, lacunar infarcts, hemorrhages, atherosclerosis, arteriosclerosis, and cerebral amyloid angiopathy^4^. A thorough description of cerebrovascular disease (CVD) assessments can be found in Wang et al^1^. Additionally, age-related neuropathological conditions, including primary age-related tauopathy (PART)^5^, aging-related tau astrogliopathy (ARTAG)^6^, and limbic-predominant age-related TDP-43 encephalopathy (LATE)^7^, were diagnosed based on standardized evaluation protocols.

To validate our diagnoses, we collaborated with the Netherlands Brain Bank (NBB), conducting a comparative staining analysis between their specimens and our brain banks. The findings demonstrated complete concordance with the diagnoses established by the NBB, reinforcing the accuracy of our assessments.

**Supplementary Figures**

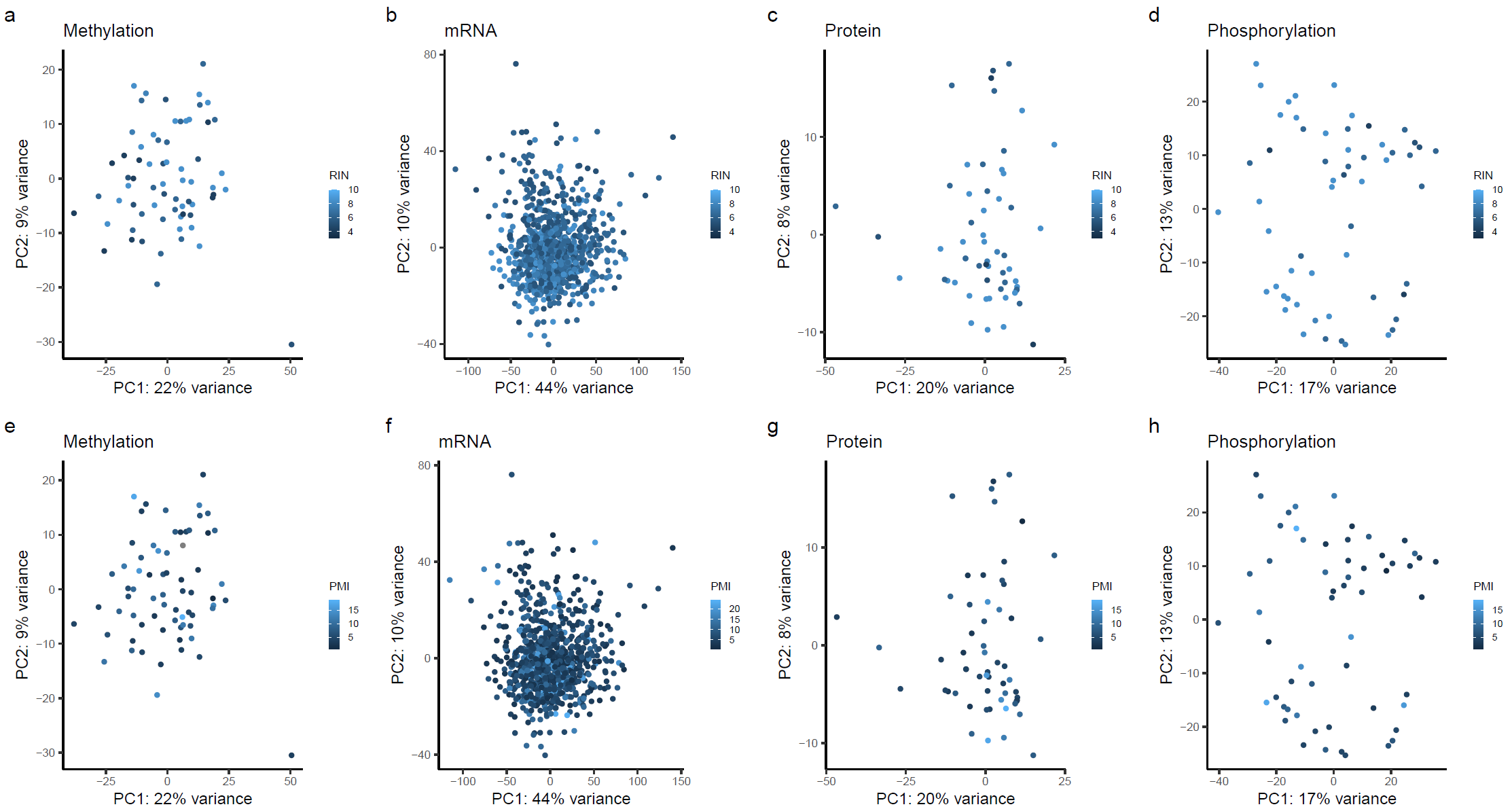

**Figure S1. Molecular profiles and sample quality features including RNA Integrity (RIN) and postmortem Interval (PMI).** Principal component analysis (PCA) was performed on methylation, mRNA expression, protein, and phosphorylation profiles of the CBMAP samples in the pilot stage. We illustrate the relationships between RIN (panels a-d) and PMI (panels e-h) with the first two principal components (PC1 and PC2) of these molecular profiles.

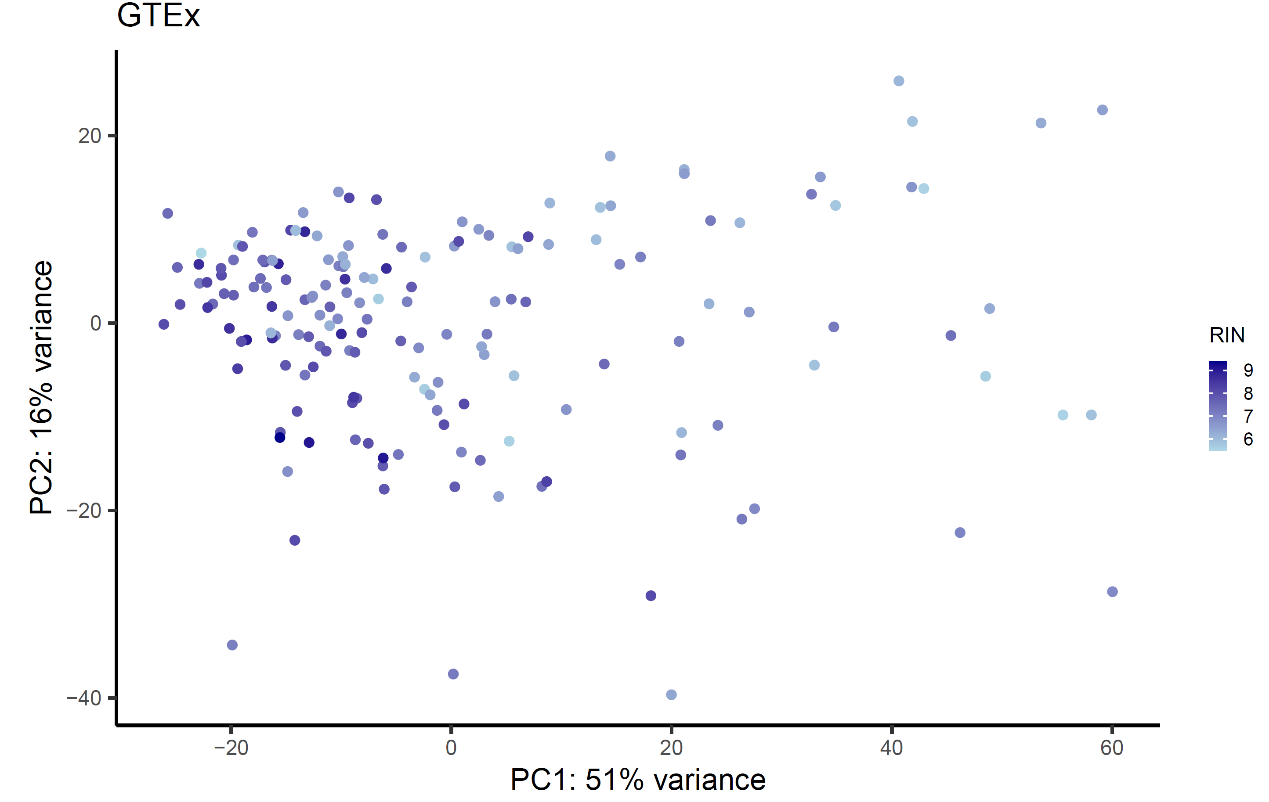

**Figure S2. A scatter plot of PC1/2 of the transcriptome profile of GTEx v8 brain frontal cortex BA9 samples.** Samples with a lower RIN and a higher RIN are colored light and dark, respectively.

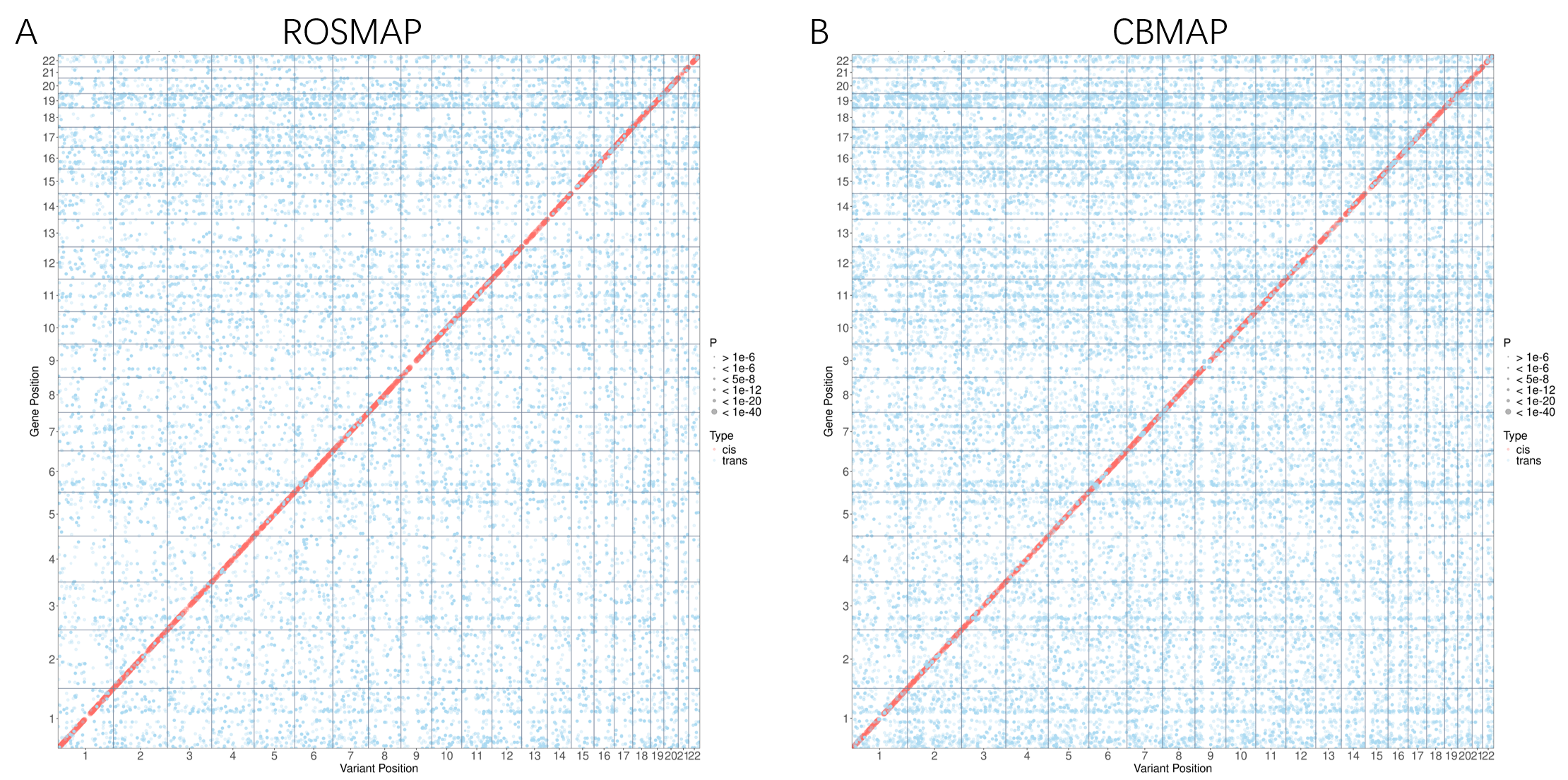

**Figure S3. Cis- and trans-eQTL signals estimated from ROSMAP and CBMAP.** QTLtools was used the estimate a linear association between SNP and gene expression by adjusting age, sex, top PCs, and batch. Red and bule dots denote cis- and trans-eQTLs. The cis region is defined by 1 Mb both sides of a gene body.

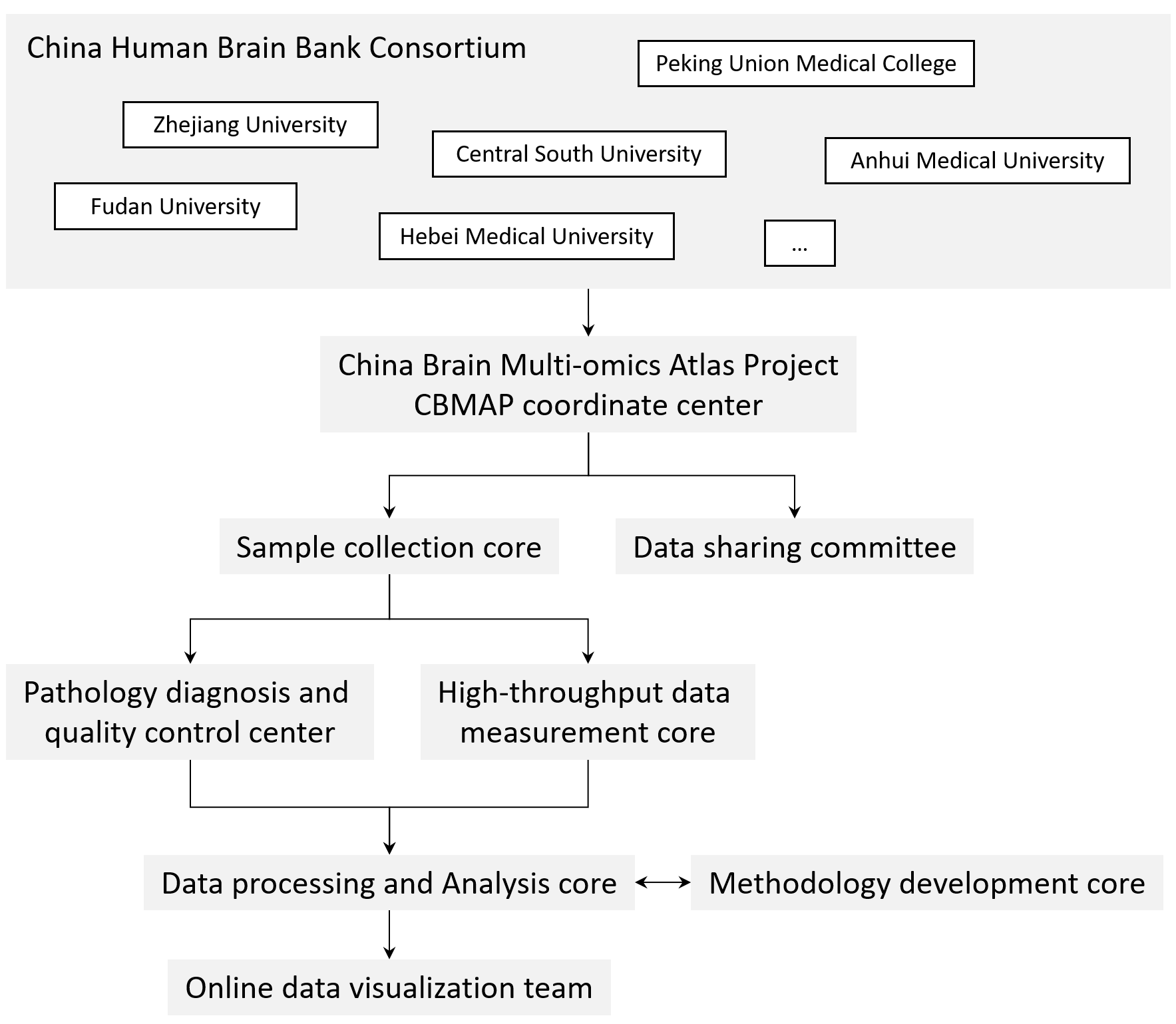

**Figure S4. Organizational Structure of the CBMAP.** The CBMAP samples are sourced from the China National Brain Bank Alliance. The coordinating center facilitates collaboration among the brain banks and various teams involved in sample collection, pathological diagnosis, high-throughput molecular assays, data processing, method development, visualization, and the Data Sharing Committee.

**Supplementary Figures**

**Table S1. Comparison among brain omics projects.**

| Project | Sample size | Ancestry | Genome | Epigenome | Transcriptome | Proteome | Metabolome | Main focus |
| --- | --- | --- | --- | --- | --- | --- | --- | --- |
| GTEx (Genotype-Tissue Expression project) | 114-237 | predominantly European ancestry | WGS | NA | mRNA | NA | NA | Understanding how genetic variation influences gene expression across multiple human tissues. |
| ROSMAP (Religious Orders Study/Memory and Aging Project) | ROS (~1,200); MAP (~1,600) | predominantly European ancestry | chip/ WGS | DNA methylation, ATAC, and histone modifications | mRNA and snRNA | Proteome (TMT) | targeted p180 metabolomics kit and Metabolon | To investigate the molecular and cellular mechanisms underlying Alzheimer's disease through multi-omics data analysis from brain tissue samples. |
| Mount Sinai/JJ Peters VA Medical Center Brain Bank (MSBB–Mount Sinai NIH Neurobiobank) | 2,000+ | predominantly European ancestry | WGS/WES | DNA methylation and ATAC | mRNA | Proteome (label-free and TMT) | NA | Integrating multiple omics data across various brain regions alongside quantitative measures of neuritic plaque density and clinical dementia ratings to advance understanding of Alzheimer's disease pathogenesis. |
| psychENCODE (phase i) | 1,866 | predominantly European ancestry | chip/ WGS | DNA methylation, histone modifications, ATAC, and Hi-C | mRNA and snRNA | NA | NA | Focusing on profiling transcriptomic and epigenomic landscapes across brain regions to understand human neurodevelopment and psychiatric disorders. |
| psychENCODE (phase ii) | 10-388 | predominantly European ancestry | WGS | snATAC-seq; single cell methylomics | snRNA and spatial transcriptomics | NA | NA | Deciphering the molecular bases of psychiatric disorders by integrating multi-omics data from human brain tissues with specific focus on single-cell and spatial transcriptomics. |
| The Charles F. And Joanne Knight Alzheimer Disease Research Center (Knight ADRC) biobank | 2,117 | predominantly European ancestry | Chip/WES/WGS | DNA methylation | mRNA and sRNA | Proteome (aptamer) | Metabolomics and lipidomics | Enhancing the understanding of Alzheimer Disease by conducting comprehensive molecular profiling across various biological samples including brain, cerebrospinal fluid, and blood. |
| CBMAP (China Brain Multi-omics Atlas Project) | 1,187 (Phase I)  2,000+  (Phase II) | **Chinese** | WGS | DNA methylation; snATAC-seq; **3D genomics** | mRNA and lncR; snRNA-seq; **Spatial transcriptomics** | Proteome (DIA); **Phosphorylation; Ubiquitination; Acetylation; etc.** | Widely targeted and targeted metabolome | Establishing a multi-omics atlas for the Chinese brain to reveal molecular regulatory networks and analyze molecular characteristics of aging processes, neurological, and psychiatric diseases. |

**Reference**

1.	Wang X, Zhu K, Wu W, Zhou D, Lu H, Du J *et al.* Prevalence of mixed neuropathologies in age-related neurodegenerative diseases: A community-based autopsy study in China. *Alzheimers Dement* 2024.

2.	Montine TJ, Phelps CH, Beach TG, Bigio EH, Cairns NJ, Dickson DW *et al.* National Institute on Aging-Alzheimer's Association guidelines for the neuropathologic assessment of Alzheimer's disease: a practical approach. *Acta Neuropathol* 2012; **123**(1)**:** 1-11.

3.	McKeith IG, Boeve BF, Dickson DW, Halliday G, Taylor JP, Weintraub D *et al.* Diagnosis and management of dementia with Lewy bodies: Fourth consensus report of the DLB Consortium. *Neurology* 2017; **89**(1)**:** 88-100.

4.	Skrobot OA, Attems J, Esiri M, Hortobágyi T, Ironside JW, Kalaria RN *et al.* Vascular cognitive impairment neuropathology guidelines (VCING): the contribution of cerebrovascular pathology to cognitive impairment. *Brain* 2016; **139**(11)**:** 2957-2969.

5.	Crary JF, Trojanowski JQ, Schneider JA, Abisambra JF, Abner EL, Alafuzoff I *et al.* Primary age-related tauopathy (PART): a common pathology associated with human aging. *Acta Neuropathol* 2014; **128**(6)**:** 755-766.

6.	Kovacs GG, Ferrer I, Grinberg LT, Alafuzoff I, Attems J, Budka H *et al.* Aging-related tau astrogliopathy (ARTAG): harmonized evaluation strategy. *Acta Neuropathol* 2016; **131**(1)**:** 87-102.

7.	Nelson PT, Dickson DW, Trojanowski JQ, Jack CR, Boyle PA, Arfanakis K *et al.* Limbic-predominant age-related TDP-43 encephalopathy (LATE): consensus working group report. *Brain* 2019; **142**(6)**:** 1503-1527.
